# On the motion of spikes: a model of multifractality as observed in the neuronal activity of the human basal ganglia

**DOI:** 10.1101/223164

**Authors:** Daniela Sabrina Andres

**Affiliations:** Laboratory of Neuroengineering, Science and Technology School, National University of San Martin (UNSAM), 25 de Mayo y Francia, 1650, San Martin, Buenos Aires, Argentina

**Author notes:** **Correspondence:** Daniela S. Andres.

**Keywords:** structure function, multifractal spectrum, globus pallidus interna, Parkinson’s disease, basal ganglia, neural field, velocity field, interspike intervals, nonlinear differential equations, diffusion coefficient

## Abstract

Neuronal signals are usually characterized in terms of their discharge rate. However, this description is inadequate to account for the complex temporal organization of spike trains. In particular multifractality is a hallmark of the neuronal activity of the human, parkinsonian basal ganglia, which is not accounted for in most models. Here I develop a new conceptualization of neuronal activity, enabling the analysis of spike trains in terms of a velocity field. Firstly, I show that structure functions of increasing order can be used to recover the multifractal spectrum of spike trains obtained from the globus pallidus interna (GPi) of patients with Parkinson’s disease. Further, I propose a neural field model to study the observed multifractality. The model describes the motion of spikes in terms of a velocity field, including a diffusive term to consider the physical properties of the electric field that is associated to neuronal activity. As the model is perturbed with colored noise, the following is observed: 1. multifractality is present for a wide range of diffusion coefficients; and 2. multifractal temporal properties are mirrored into space. These results predict that passive electric properties of neuronal activity are far more relevant to the human brain than what has been usually considered.

## 1 Introduction

The basal ganglia are a circuit of densely interconnected subcortical nuclei, whose disease is related to human movement disorders (Obeso, Rodriguez-Oroz et al. 2008). These disorders include Parkinson's disease, the second most common neurodegenerative pathology following Alzheimer’s disease (Kleinman and Frank 2013). Due to the aging of the population, the social burden of neurodegenerative disorders is increasing worldwide, deepening the need to understand their pathophysiology. Current models of the basal ganglia are partly successful in the prediction of neurophysiologic alterations occurring in movement disorders. However, no current model allows to predict and control deep brain stimulation (DBS), one of the major therapeutic approaches to Parkinson’s disease (Montgomery Jr 2016). The lack of a description of the complex properties of the basal ganglia neuronal activity might be one cause of this failure. The fundament of the classic model of pathophysiology of the basal ganglia lies on a description of the discharge rate of their output nuclei: the globus pallidus interna (GPi), and substantia nigra reticulata (SNr) (Albin, Young et al. 1989, DeLong 1990). In primates, GPi and SNr neurons fire in a tonic manner, keeping the motor thalamus inhibited, and momentary reductions of their discharge rate facilitate movement. In humans, movement onset and velocity are positively correlated to the rate of discharge of the motor thalamus, and negatively to the rate of discharge of the GPi, and SNr (Utter and Basso 2008). These observations seem to confirm the rate hypothesis, but evidence obtained during functional neurosurgery in patients with Parkinson’s disease is contradictory (for a detailed discussion see (Andres and Darbin 2017) and references therein). The predictions of the classic model of the basal ganglia are inconsistent with the effects of DBS and ablative surgery, raising concerns on the validity of the rate of discharge as a measure of the state of the basal ganglia (Tang, Moro et al. 2005, Nambu and Chiken 2015).

Nonlinear time series analysis has inspired an alternative view. A growing body of evidence indicates that complex properties are crucial for the understanding and modeling of basal ganglia pathophysiology (Obeso, Rodriguez-Oroz et al. 2000, Darbin, Soares et al. 2006, Lim, Sanghera et al. 2010, Alam, Sanghera et al. 2015). Neuronal spike trains of the basal ganglia are commonly recorded during DBS surgery with clinical purposes; statistical, frequential, and nonlinear methods have been extensively applied to their study (Levy, Dostrovsky et al. 2001, Brown 2003, Hutchison, Dostrovsky et al. 2004, Alonso-Frech, Zamarbide et al. 2006, Lee, Verhagen Metman et al. 2007, Gale, Amirnovin et al. 2008, Hohlefeld, Ehlen et al. 2013). Yet, their characterization is not complete. In previous work we analyzed first order temporal structure functions of basal ganglia spike trains, and identified non-random time patterns (Andres, Cerquetti et al. 2015, Nanni and Andres 2017). Structure functions were developed initially to describe the behavior of turbulent fields in the inertial range, and have been later applied to the analysis of physiologic signals (Kolmogorov 1941, Muzy, Bacry et al. 1993). Interestingly, structure functions exhibit scaling properties that allow classifying a signal as monofractal or multifractal (Hayot and Jayaprakash 1996, Lin and Hughson 2001). To that goal, temporal structure functions of order higher than one are calculated now from GPi spike trains, recorded during DBS surgery. Results indicate that a multifractal spectrum is present in the neuronal discharge of the GPi from patients with Parkinson's disease. This complex geometric behavior can be interpreted as a hallmark of the human, parkinsonian basal ganglia. Since the GPi is one of the main output nuclei of the basal ganglia, the results obtained from GPi spike trains are representative of the processing of information performed by the whole circuit.

The arguments presented suggest that considering multifractality in mathematical models of the basal ganglia is likely to be of fundamental importance to achieve accuracy. Here, multifractality is reproduced using a neural field to model the velocity of spikes as they move through a neural network. The model consists of a nonlinear differential equation that includes a gradient term and a diffusive term. Diffusive features are present in neuronal processes, given the physical properties of electric fields. However, the role of these so-called passive electric properties of neuronal activity is not usually considered in neurophysiology studies, which more commonly focus solely on spiking activity. Previous modeling work on the basal ganglia by our group showed that a diffusion coefficient is a critical parameter for the control of information transmission and information deterioration in the parkinsonian GPi (Andres, Gomez et al. 2014). The model introduced here shows that: 1. multifractality of neuronal spike trains is related to diffusive properties, and 2. multifractality of temporal activity is reflected on the spatial domain. These results predict that passive (diffusive) properties of neuronal activity determine the structure of temporal and spatial neuronal activity in the basal ganglia, and must be considered in the study and treatment of movement disorders.

## 2 Methods

### 2.1 Patients and signal recording

Six patients fulfilling the clinical criteria of the United Kingdom’s Parkinson’s Disease Society Brain Bank for idiopathic Parkinson’s (UK-PDS-BB) disease, Hoehn & Yahr IV, underwent stereotactic neurosurgery and were included in this study (Hoehn and Yahr 1967). All patients were on chronic treatment with L-DOPA, presented similar motor affectation, with severe dyskinesias and motor fluctuations, and fulfilled the Core Assessment Program for Surgical Interventional Therapies in Parkinson’s Disease (CAPSIT) criteria for inclusion in the surgery program (Defer, Widner et al. 1999). All patients signed informed consent prior to the procedure, previously reviewed and approved by the institutional ethics committee, and were without medication at the time of the surgery and during data acquisition. Age, gender and other details are not shown to protect patients’ privacy.

Details about the recording procedure have been already published elsewhere (Andres, Cerquetti et al. 2011). Briefly, microrecording, stimulation and neurosurgical procedures were performed in patients awake, under local anesthesia. Surgical targets were planned employing magnetic resonance imaging (MRI) and using a Leksell stereotactic system (Series G, Elekta, Sweden). Microrecordings of neuronal activity were obtained only during the surgery, after the GPi was identified by an expert. Platinum/iridium (Pt/Ir 80/20%) microelectrodes with nominal impedance of 0.8-1.2 megohms (mTSPBN-LX1, FHC Inc) were used. A differential amplifier with a built-in impedance meter (FHC IS-AM-00-01 Iso-Xcell 3 Amplifier) and an isolated stimulus generator (FHC IS-PL-06 Isolated Bipolar Pulsar Stimulator; FHC, Bowdoinham, ME) were connected to a preamplifier remote probe mounted onto a motorized microdrive (FHC 65-00-1 Stepper Drive and ST-M0-00 TMS Controller), located near the electrode tip to minimize pickup of electrical noise. A dedicated acquisition system (1401plus, CED) was employed to amplify, and digitize the signal, filtering with second order 300 to 5,000 Hz bandpass and 50 Hz line notch analog filters. The sampling rate was 50 kHz. The total amplification including probe was set at gain ×10000 (checked with a built-in calibration signal of 1mV at the beginning of each surgery).

### 2.2 Data analysis

Signals were processed off-line. To obtain single cell recordings, spikes were extracted from raw signals and separated into classes using wavelet analysis and clustering (Quiroga, Nadasdy et al. 2004). From these single cell data, time series of interspike intervals (ISI) were constructed. Following that, the temporal structure function was computed as follows. An interval I(t) is defined as the *t*^th^ ISI, which is used to calculate the increment ΔI(τ) = *I*(t + τ) − *I*(t), *τ* being the time lag between 1 and 1000. The temporal structure function S_q_ (τ) is

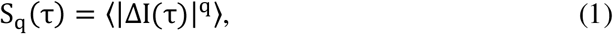

where 〈·〉 is the statistical average, and q is the order of the structure function, an integer between 1 and 30. Next, S_q_ was smoothed by applying a running average over a 30 points window to obtain a smoothed structure function, 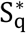 (30 points shorter than S_q_). The behavior of interest here is the scaling behavior of 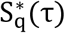. Given the general relationship

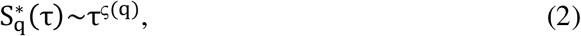

the scaling exponent is a function, ς_τ_(q), which defines the fractal properties of the signal under study. For random processes, *ς*_τ_(q) = 0. In the case of monofractals the exponent function grows linearly, ς_τ_(q) = qς(1). If the signal is multifractal, ς_τ_(q) is a nonlinear function (Hayot and Jayaprakash 1996, Lin and Hughson 2001)^(1)^.

To find ς_τ_(q), 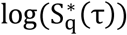 vs. log(τ) was considered, and a linear (scaling) region was looked for. The scaling region was defined as the longest range of τ fitting a linear function with non-zero slope (ς≠0), for at least 1 ≤ q ≤ 10. Linear regressions were considered acceptable if R^2^ ≥ 0.6, and regressions with regression coefficients smaller than that were discarded.

### 2.3 Model and simulations

This model describes neuronal activity in terms of the velocity of spikes. While spikes travel with constant velocity along axons, the same does not hold for neural tissue or neural networks at larger scales. Indeed, consider the following simplified situation. Take a single spike *S* that arrives to a neuronal soma whose membrane potential *V* is below the triggering threshold 𝜗. If this spike induces an excitation with an amplitude *e* in the neuron, such that the *V* + *e* > 𝜗, then the neuron will fire in response, which is equivalent to saying that the spike *S* has travelled through the neuron (a node of the network). Importantly, the condition *V* + *e* > 𝜗 can be satisfied for a range of potentials *V* and, since summation is a time-consuming mechanism, the spike triggered in response to the excitation *e* will be fired faster for membrane potentials that are closer to the threshold, giving place to a higher velocity of transmission *u*. Not only is velocity variable, but it is nonlinearly so. The process of summation is typically a nonlinear function of time, introducing nonlinear variability in the profile of *u*. Although the velocity of spikes *u* is not a common variable in neurophysiologic studies, the related variable of neuronal frequency *f* is present in a large part of the literature. The relation between both variables is a simple one: since *u* = Δ*x*/Δ*t* and *f* = 1/Δ*t*, then *u* = *f* · Δ*x*. Consequently, for distances Δ*x* = 1 the module of the velocity |*u*| has the same value than the frequency. Regarding the time intervals between spikes (interspike intervals *I*, ISI), they are equal to the inverse of the instantaneous frequency of discharge, and therefore |*u*| = 1/*I*. Due to this relation, it is expected that any nonlinearity of *u* should be reflected on the time series of interspike intervals.

To model the behavior observed in the clinical data, the following neural field equation is introduced:

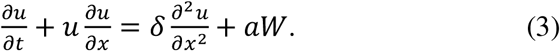

Here, *u*(*x*, *t*) represents the velocity of spikes (a function of time *t* and space *x*), *δ* is a diffusion coefficient and *W* is a stochastic drive, described in more detail below. Note that this model takes the form of stochastic Burgers’ equation, which has been intensively studied in the field of fluid dynamics (Burns, Balogh et al. 1998). In previous work I proposed that this kind of equations might be useful to study mathematical properties of neuronal activity, which is further analyzed here (Andres, Cerquetti et al. 2011).

Computer simulations were run with custom Matlab and R code. The model was studied numerically in 1D. The discretization scheme selected was a Crank-Nicolson scheme, which is suitable for this kind of study (Kadalbajoo and Awasthi 2006). The space steps and time steps were Δ*x* = Δ*t* = 0.001, and *u*(*t*) = 0 at the boundaries of the spatial domain. The initial condition considered was a random distribution of *u*(*x*) between −1 and 1, with a power spectrum obeying the scaling law *P*(*ω*(*u*)) = *ω^β^*, where *β* < 0. Following Hayot et al., the same kind of colored noise *W* was used as a drive to the equation, with an amplitude *a* = 10 ^−6^ (Hayot and Jayaprakash 1996). The temporal evolution of *u*(*x*) was studied for 1000 time and space steps and for a range of diffusion coefficients 10^−3^ < *δ* < 0.11, varying *δ* in increments of 0.02. To make the data obtained from the model more directly comparable to clinical data, the inverse of the absolute value of the velocity was considered, obtaining a variable equivalent to *I*(*x*,*t*). This variable 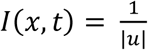 was analyzed applying the same algorithm that was described in point 2.2 for the analysis of clinical ISI time series to obtain a temporal structure function *S_q_*(τ) at fixed points in space. Finally, a spatial structure function *S_q_*(*X*) was computed in a similar manner, but considering adjacent points of *u* for fixed times. Scales between 1 and 50 were used to compute the structure functions both in space and time. A multifractal spectrum was looked for both in the space (ς_*X*_(q)) and time domains (ς_τ_(q)), by analyzing the behavior of *S*_q_(τ)~τ^ς(q)^ and *S*_q_(X)~X^ς(q)^ for *q* varying between 0 and 2 in steps of 0.1.

## 3 Results

Results for the analysis of clinical data are shown in figures-1 and 2. In total, 22 GPi recordings obtained from 6 patients were analyzed. Time series had a mean length of 5668 ± 773 ISI (mean ± standard error of the mean, SEM). A scaling region of the temporal structure function was found for a minimum range of 1 ≤ *q* ≤ 10 in every neuron. In 15 neurons, the fitting was sufficiently good for q up to 30 (*q_max_* = 24 ± 2, mean ± SEM). The length of the scaling interval over *τ* was 79 ± 25; mean ± SEM. Figure 1 shows smoothed structure functions of order 1 to 6 and the corresponding scaling region for a sample neuron. In figure 2 the function ς_τ_(q) for sample neurons is shown. In 18 cases (82%) a nonlinear ς_τ_(q) was found, implying a multifractal organization of the spike trains analyzed.

**Figure 1).**
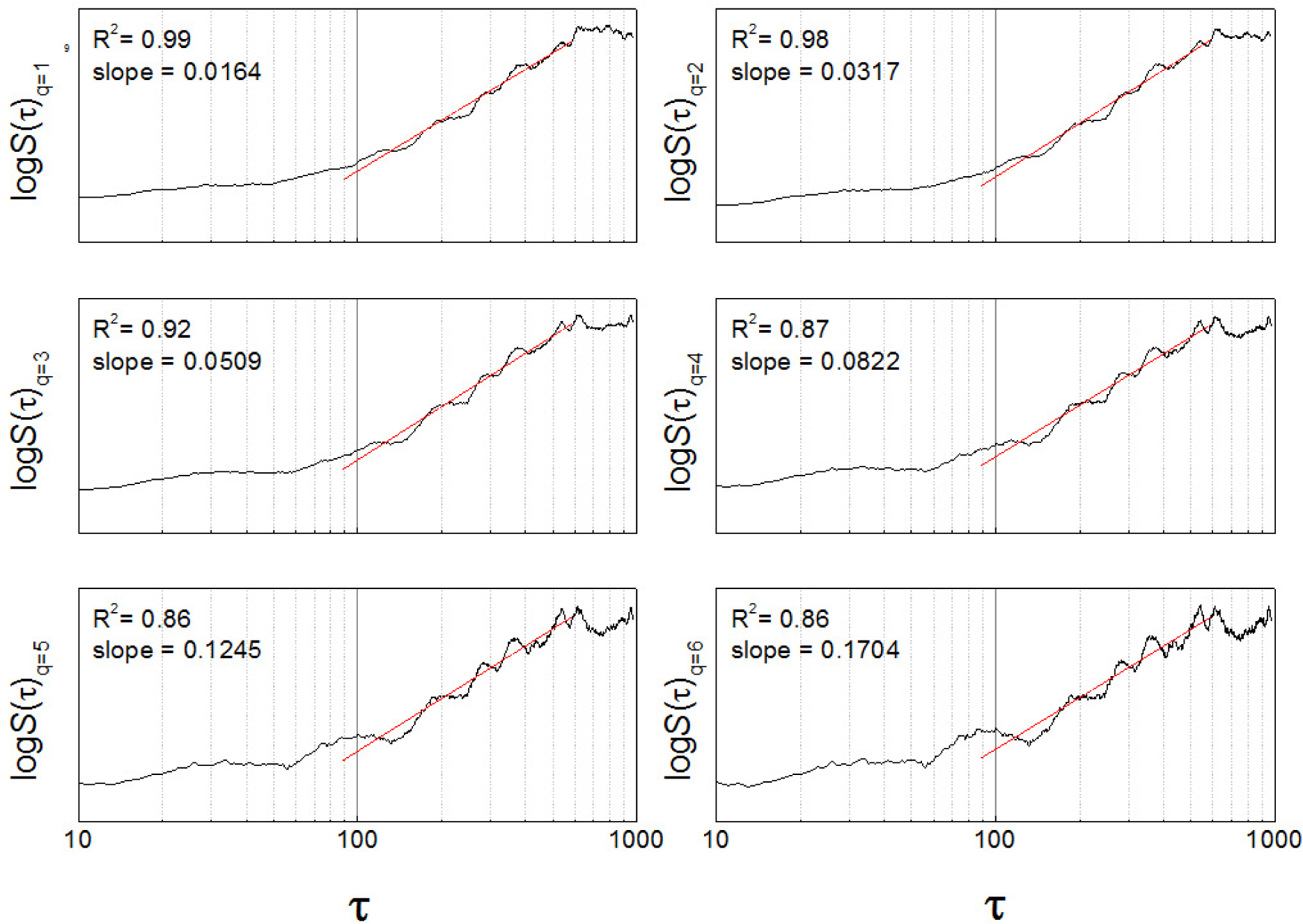
Structure functions of order 1 to 6 for a sample neuron, showing the scaling region (89<τ<589) and corresponding linear fittings.

**Figure 2).**
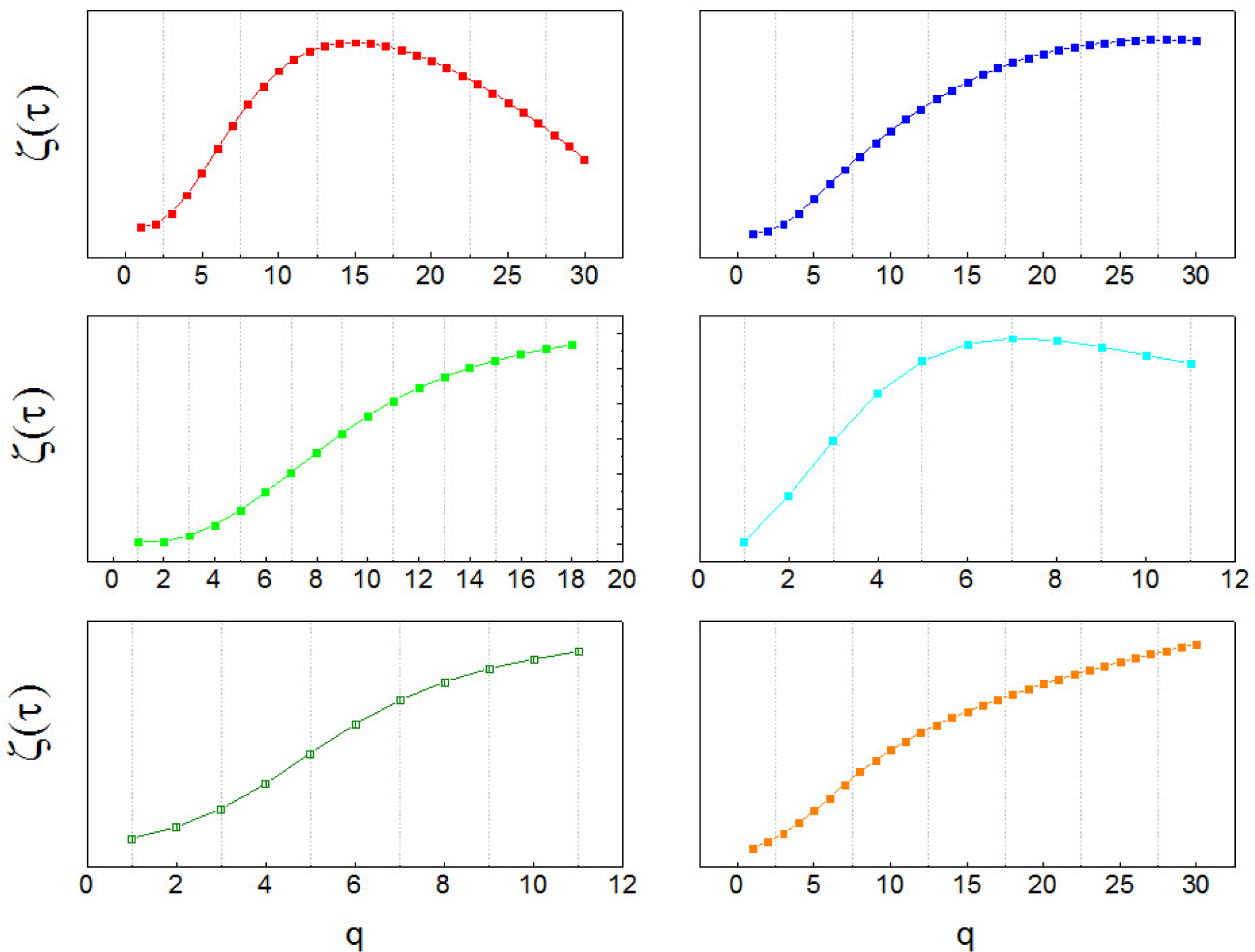
Multifractal spectrum of sample neuronal recordings. The function *ς*_τ_(*q*) is built as the slope of the scaling region in log *S_q_*(*τ*), vs. the order in logarithmic scale, *q*.

Figure 3 shows the results of computer simulations. On the left column, the white areas represent the areas where 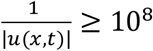, and the black areas correspond to the parts of the domain where the module of the velocity of spikes (equivalent to frequency) is highest. As the diffusion coefficient increases (figure 3, from top to bottom), the white areas are enlarged, and the total area of higher velocity diminishes. However, at the same time the whole integration domain turns less heterogeneous, with individual areas of high velocity becoming wider. This indicates that diffusion not only contributes to the dissipation of energy (lowering the global velocity of spikes transmission), but it also modifies the temporal and spatial organization of the neuronal activity across the neural field. Importantly, hallmarks of multifractality appear both in space and time for the whole range of *δ* investigated. The middle and right columns of figure 3 show typical examples of ς_*τ*_(q) and ς_*X*_(q), respectively. Observe that multifractality is characteristic of the model, with nonlinear exponent functions in the spatial as well as temporal domains.

**Figure 3).**
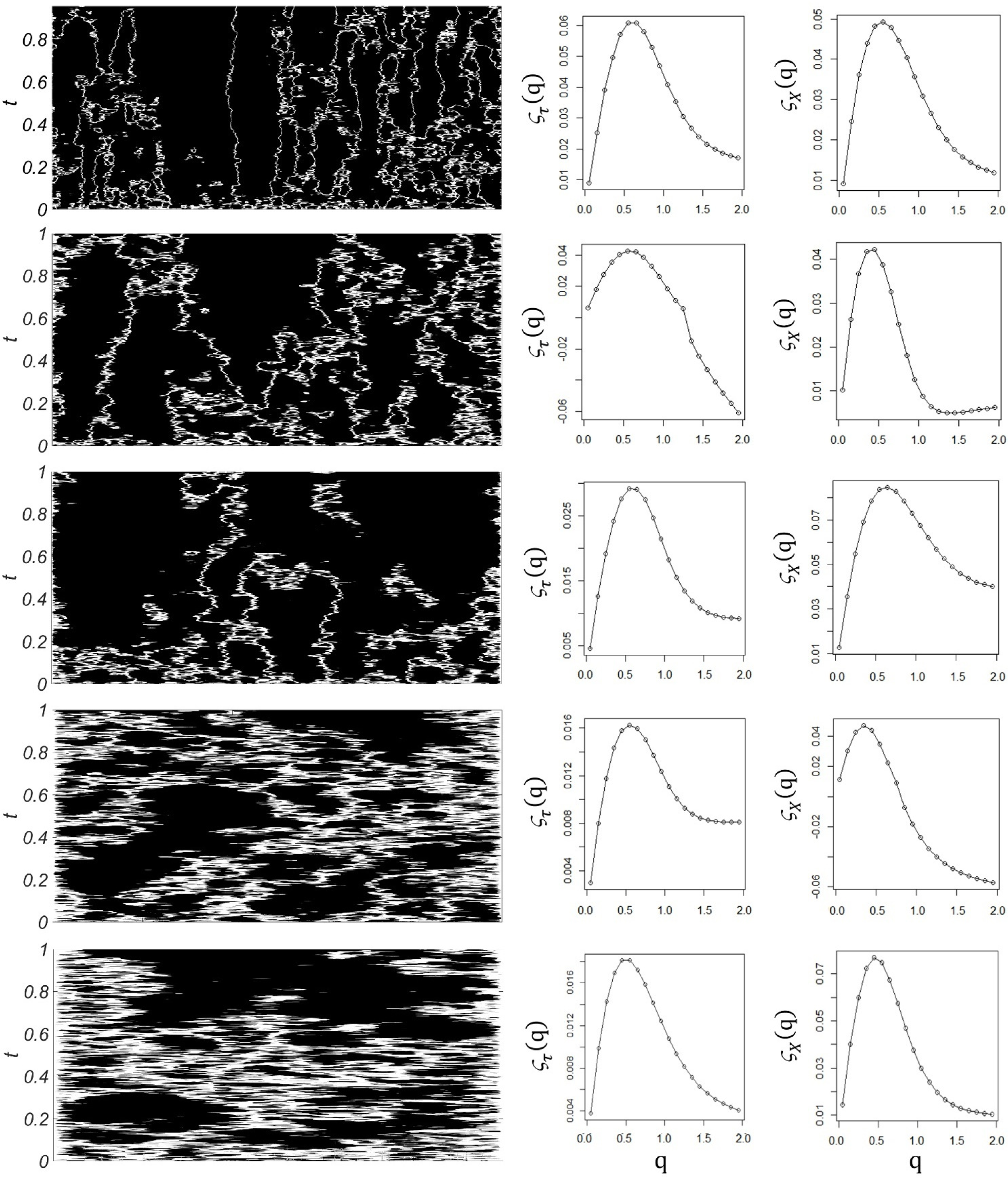
Left column: Velocity field *u*(*x*,*t*), where *x* is space and *t* is time. Middle column: Temporal exponent function, *ς*_τ_(*q*). Right column: Spatial exponent function, *ς_X_*(*q*). The diffusion coefficient *δ* increases from top to bottom (0.001; 0.005; 0.01; 0.05; 0.11). On the left column, white areas represent the parts of the domain where 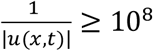, while in black areas the velocity is higher. As *δ* grows, the areas of high velocity become wider, but the total are of high velocity diminishes. Nonlinearity governs the exponent function *ς*(*q*) in the space as well as in the time domains, indicating multifractality.

## 4 Discussion and conclusion

The focus of this paper is on the structure of spike trains produced by GPi neurons of the human, parkinsonian basal ganglia, and what might be a suitable mathematical model of it. Lately, it is getting clear that the discharge rate is insufficient to characterize the neuronal activity of the basal ganglia. This approach fails, because it doesn’t predict the response of the basal ganglia to DBS, a highly effective therapy to ameliorate parkinsonian symptoms (Loher, Burgunder et al. 2002, Wichmann and Delong 2006, Sestini, Pupi et al. 2007, Obeso, Rodriguez-Oroz et al. 2008, Eusebio, Thevathasan et al. 2011, Stefani, Fedele et al. 2011, Miocinovic, Somayajula et al. 2013, Chiken and Nambu 2014). This has motivated the research of other, quantitative features of neuronal activity as candidate measures of the state of the system. In particular, several authors offered evidence of complex time patterns in the neuronal activity of the basal ganglia (see for example the discussion in (Nambu 2005)). In previous work I analyzed first order structure functions (*q* = 1) of simulated, animal, and human spike trains, and hypothesized that time patterns and rate-coding properties co-exist in the basal ganglia (Andres, Gomez et al. 2014, Andres, Cerquetti et al. 2015, Andres, Cerquetti et al. 2016). Here I continue the study to higher orders (*q* > 1) and show that the scaling behavior of structure functions can be used to obtain the multifractal spectrum of the human, parkinsonian basal ganglia. This level of description is relevant for translational studies, which connect various models of Parkinson's disease to the clinic, since data obtained during functional neurosurgery offer a unique opportunity to study the human, parkinsonian basal ganglia directly. Mathematical descriptions of complex properties of the human basal ganglia lay the foundation for basic and translational research on Parkinson's disease. The algorithm employed is robust to short, noisy time series and efficient computationally, showing potential for on-line, neurosurgical applications. Alternative methods of multifractal analysis include multifractal detrended fluctuation analysis and wavelet-based methods (Muzy, Bacry et al. 1993, Peng, Havlin et al. 1995, Stanley, Amaral et al. 1999). A comparison with other methods remains for future studies.

The clinical findings of this work agree with the literature. Zheng et al. used a multiplicative model to reproduce fractal features of human, basal ganglia neuronal activity (Zheng, Gao et al. 2005, Rasouli, Rasouli et al. 2006). Other complex and nonlinear temporal properties are also present in basal ganglia spike trains from rodents, primates and patients with movement disorders (Li, Jia et al. 2008, Lim, Sanghera et al. 2010, Hohlefeld, Huebl et al. 2012, Darbin, Adams et al. 2013, Hohlefeld, Ehlen et al. 2015). The most popular nonlinear measure for the characterization of parkinsonian spike trains is probably entropy, which diminishes due to pharmacological or stimulation treatment, and increases due to alertness (Latora 2000, Lafreniere-Roula, Darbin et al. 2010, Lim, Sanghera et al. 2010, Andres, Cerquetti et al. 2014, Alam, Sanghera et al. 2015, Darbin, Jin et al. 2015). Importantly, entropy can be estimated wrongly if the multifractality of a system is not considered (Costa, Lyra et al. 1997, Lyra 1998, Latora 2000).

Finally, this work might have an impact not only on movement disorders, but also on other areas of neuroscience. Considering spike trains from the perspective of their velocity introduces a new concept to the analysis of spiking neurons. As was discussed before, interspike intervals (a measure commonly used in neurophysiology studies) can be regarded as the inverse of the absolute value of the velocity of spikes. Following this idea, neural field models can be used to model neuronal activity as a velocity field, *u*(*x*,*t*). Here, a nonlinear differential equation is introduced that considers *u*(*x*,*t*) as a combination of a nonlinear gradient term and a diffusive term. This is relevant to Parkinson’s disease, where it has been hypothesized that increased diffusion plays a critical role in the deterioration of information transmission (Andres, Gomez et al. 2014). In this context, a progression of Parkinson’s disease would be toward higher diffusion coefficients, lowering the velocity of spikes and homogenizing the velocity field. Nevertheless, the model produces complex geometric structures like the ones observed in clinical data for the whole range of coefficients studied. These complex structures are mirrored from time into space: a multifractal spectrum is observed in both the temporal and the spatial domains. In previous work I pointed out that there are mathematical similarities between the neuronal activity of the parkinsonian basal ganglia and turbulence (Andres, Cerquetti et al. 2011, Andres, Gomez et al. 2014). Now I introduce a neural field model inspired by Burgers' equation, a well-known equation in the field of fluid dynamics. The model is based on some physical properties of neurons, and it captures essential features of neuronal activity of the human brain, reproducing multifractality as observed in the neuronal activity of the human, parkinsonian basal ganglia.

## Footnotes

(1) Different authors use different notations. The function that we call ς_τ_(q) is called τ(q) in Touchette’s work, according to whom the spectrum of singularities *f*(*α*) does not need to be concave (Touchette and Beck 2006). A concave *f*(*α*) can be calculated as the Legendre transform of ς_τ_(q), whereas a not concave *f*(*α*) cannot. In any case, a concave ς_τ_(q) implies multifractality.

## 6 Author Contributions

DSA: study design, data analysis, computer simulations, manuscript preparation

## 8 Acknowledgments

The author kindly acknowledge the work of all the medical and technical personnel at Fleni Institute, who made this work possible.

